# Loss of resistance to punishment of cocaine use after prior experience

**DOI:** 10.1101/2020.07.24.219170

**Authors:** Audrey Durand, Paul Girardeau, Luana Freese, Serge H. Ahmed

**Author notes:** **Correspondence to**: Serge H. Ahmed, Ph.D., Université de Bordeaux, Institut des Maladies Neurodégénératives, UMR CNRS 5293, 146 rue Léo Saignât, 33076 Bordeaux, France.

## Abstract

One behavioral feature of drug addiction is continued drug use despite awareness that this causes negative consequences. Attempts to model this feature in animals typically involve punishing drug self-administration with a brief electrical footshock and look for resistance to punishment. Though all individual animals eventually stop self-administering the drug with increasing intensity of punishment, some individuals do so at higher intensities than other individuals. The greater relative resistance to punishment of the former individuals is generally interpreted as evidence for a compulsion-like behavior. Here we show that resistance to footshock punishment is in fact not a stable individual behavioral feature. Specifically, when rats are retested for their resistance to increasing intensity of footshock punishment, they become much less resistant. As a result, they suppress their cocaine intake even when punished with an initially low and ineffective intensity. A series of original behavioral experiments reveals that this low resistance to footshock punishment is rapidly acquired after rats experience a punishment intensity that leads them to near-completely suppress their cocaine intake. Passive exposure to the same intensity does not induce this effect. Once acquired, low resistance to punishment persists during at least one month, but can nevertheless be extinguished by retesting rats on a daily basis. Interestingly, this acquired low resistance to footshock punishment does not generalize to a non-painful form of punishment (i.e., histamine) that is also seldom used in animal drug self-administration studies. We discuss some possible theoretical and methodological implications of these findings for future research on animal models of compulsion-like behavior.

## Introduction

One behavioral feature of drug addiction is continued drug use despite awareness that this causes negative consequences [1–3]. Though becoming aware of a causal relationship between drug use and negative consequences can be difficult, when such awareness eventually emerges it often motivates affected individuals to try quitting drug use, even if these attempts often fail, at least initially, and end up in relapse [2,4,5]. It is precisely when individuals attempt to quit drug use to avoid the associated negative consequences, but with no success, that a compulsion-like state is typically inferred [1,6,7]. Thus, confirming that individual drug users have developed addiction requires at least three conditions: i) drug use causes negative consequences, ii) drug users are aware of this fact and iii) they have attempted to quit several times and have repeatedly failed – at least initially [1]. The latter clause is added because after several unsuccessful attempts, many drug users eventually succeed to quit drugs [8].

Despite some initial observations [9–12], it is only recently that the role of negative consequences in defining addiction-like behavior in animal models has come under intense scrutiny [13–15]. Interestingly, since drug self-administration is studied under relatively safe laboratory conditions, it has little direct negative consequences which may contribute to explain why animals do not typically attempt voluntarily to abstain from drug use [16]. In other words, in a standard drug self-administration setting, condition i) is rarely met. Negative consequences have to be added to drug self-administration. This often involves punishing drug self-administration with a brief electrical footshock (FS) [9,17–19]. For instance, in a typical punishment experiment, after rats have learned an operant response (e.g., pressing a lever) to self-administer an intravenous drug, this response is punished by an immediate FS while it continues to be also rewarded by the drug, though not always. When the intensity of FS punishment is fixed (i.e., typically around 0.4 mA), one can observe that some animals stop using the drug while others resist and continue using the drug despite FS punishment [17,19,20]. Though we do not know whether and to what extent the latter individual animals can meet conditions ii) and iii) above [21,22], their resistance to punishment has generally been interpreted as evidence for a compulsion-like state and its associated neuronal substrates have been interpreted in this light [19,20,23,24].

Though there is clear evidence for individual variation in resistance to FS punishment, this variation is not all-or-none. Parametric studies that have tested a broad range of FS intensities present a more nuanced picture [25–37]. Below some low intensities, all individuals are resistant to FS punishment and, inversely, above some high intensities, all individuals stop using the drug. As expectable, individual variation manifests between these two extremes, with some individuals stopping drug use at higher FS intensities than other individuals. This simple observation is important because it shows that what differs between individuals is not their ability to stop drug use *per se*, since all individuals eventually do so, but their relative resistance to increasing punishment before stopping drug use. This observation may beg the question of whether and to what extent a high resistance to FS punishment is a better model of compulsion-like behavior than a low resistance. However, before addressing this question, it is important to demonstrate that such relative resistance to punishment is a stable individual feature.

The present study was designed to address this question. We tested cocaine self-administering rats with a broad, within-session range of FS intensities (i.e., 0.1-0.9 mA) to measure the intensity that suppresses by 50% the baseline rate of drug self-administration.

We used this measure as a quantitative proxy for resistance to FS punishment. After a first assessment of resistance to FS punishment, rats were allowed to recover their pre-punishment levels of drug self-administration before being retested for resistance to punishment. Our main finding shows that individual resistance to FS punishment is not stable. Specifically, during the second assessment, rats were much less resistant to punishment than during the first assessment and near-completely suppressed their cocaine intake when punished with an initially low and ineffective FS intensity. We thus conducted a series of original behavioral experiments to try to characterize this acquired low resistance to FS punishment. Overall, our findings show that prior experience with a high-intensity FS punishment makes rats less resistant to subsequent punishment.

## Methods

### Subjects

A total of 60 adult male Wistar Han rats (225-250g at the beginning of experiments; Charles River, Lyon, France) were used in this series of experiments. Rats were housed in groups of 2 or 3 and were maintained in a light-(reverse light-dark cycle), humidity-(60 ± 20%) and temperature-controlled vivarium (21 ± 2°C). All behavioral testing occurred during the dark phase of the light-dark cycle. Food and water were freely available in the home cages throughout the duration of the experiment. Home cages were enriched with a nylon gnawing bone and a cardboard tunnel (Plexx BV, The Netherlands). 22 rats did not complete the behavioral experiments which lasted several months, thereby leaving a total of 38 rats for final analysis. Rats did not complete the experiments due to a variety of factors (e.g., failure to acquire cocaine self-administration; infection; catheter failure).

### Ethical statement

All experiments were carried out in accordance with institutional and international standards of care and use of laboratory animals [UK Animals (Scientific Procedures) Act, 1986; and associated guidelines; the European Communities Council Directive (2010/63/UE, 22^th^ of September 2010) and the French Directives concerning the use of laboratory animals (Decree 2013-118, 1^st^ of February 2013)]. The animal facility has been approved by the Committee of the Veterinary Services Gironde, agreement number A33-063-922.

### Apparatus

Six identical operant chambers (30 × 40 × 36 cm) were used for all behavioral testing and training (Imetronic, Pessac, France). They were located away from the colony room in a separate dimly lit room. They were individually enclosed in sound-attenuating wooden cubicles equipped with a white noise speaker (45 ± 6 dB) for sound attenuation and an exhaust fan for ventilation. Each chamber was equipped with two retractable metal levers on opposite panels of the chamber, and a corresponding white light diode positioned above each lever. One syringe pump delivered drug solution through Tygon tubing (Cole Parmer, Vernon Hills, IL, USA) connected via a single channel liquid swivel (Lomir Biomedical Inc., Quebec, Canada) to a cannula connector (Plastics One, Roanoke, VA, USA) on the back of the animal. This system of drug self-administration was suspended at the center of the chamber. Finally, the grid floor of each chamber was connected to a generator that delivered scrambled electric footshock. The onset, duration and intensity of each footshock were programmed by the experimenter (see below).

### Surgery

Three days after their arrival in the laboratory, rats were anesthetized with Xylazine (15 mg/kg, intraperitoneal (i.p.), Merial, Lyon, France) and Ketamine (110 mg/kg, i.p., Bayer Pharma, Lyon, France) and were surgically prepared with an indwelling silastic catheter (0.012 inch inner diameter, 0.025 inch outer diameter, Dow Corning Corporation, Michigan, USA) in the right jugular vein. The catheter was secured to the vein with surgical silk sutures and passed subcutaneously to the top of the back about 2 cm below the scapulae where it exited into a connector (modified 22 gauge cannula). After surgery, animals were flushed daily with 0.2 ml of a sterile ampicillin solution (0.1 g/ml, Panpharma, Fougères, France) containing heparin (300 IU/ml) to maintain patency. When a leakage in the catheter was suspected, its patency was checked by an intravenous administration of Etomidate (0.75-1 mg/kg, Braun Medical, Boulogne-Billancourt, France), a short-acting non-barbiturate anesthetic. Behavioral procedures began 7-10 days after surgery.

### Drugs

Cocaine hydrochloride (Coopération Pharmaceutique Française, Melun, France) was dissolved in 0.9 % NaCl, filtered through a syringe filter (0.22 μm) and stored at room temperature (21 ± 2 °C). Drug doses are expressed as the weight of the salt.

### Data analysis

All data were subjected to relevant repeated measures ANOVAs, followed by Tukey *post hoc* tests where relevant. Statistical analyses were run using Statistica, version 7.1 (Statsoft Inc., Maisons-Alfort, France). In all experiments, the first 30-min interval of each session was systematically ignored in within-session analysis of behavior because it includes the initial drug loading phase. During drug loading, cocaine self-administration can indeed be much higher than during the rest of the session [38–40]. In experiment 1, the current intensity that suppresses by 50% (IS50%) the rate of cocaine self-administration was assessed by fitting each individual intensity-effect curve with a two-parameter sigmoid function (SigmaPlot 8.02, SPSS Inc., Chicago, IL).

### General behavioral procedures

#### Cocaine self-administration training

In all experiments, animals were first habituated during 2 3-h daily sessions to the operant chamber and the tethering system. During habituation, no lever was presented, and rats were allowed to move freely to explore the chamber. After habituation, rats were progressively trained to press a lever to self-administer cocaine intravenously (0.25 mg per injection) under a fixed-ratio (FR) 3 schedule of reinforcement during 4-5 weeks. All self-administration sessions began with extension of the operant lever and ended with its retraction after 2.5-4h, depending of the experiment (see below). No inactive lever was used in the present study. Intravenous delivery of cocaine began immediately after completion of the operant response requirement and lasted about 4 s. It was accompanied by illumination of the light cue above the lever for 20 s. Responses during the light cue were recorded but had no programmed consequence. All self-administration sessions were run 5 days a week.

#### Footshock punishment of cocaine self-administration

In all experiments, sessions of footshock (FS) punishment were subdivided into 2 main periods: a first period of cocaine self-administration without punishment that lasted 1h followed by a second period where cocaine self-administration was punished (see Figure 1). The first period served as a control to measure the effects of FS punishment on cocaine self-administration during the second period. During the latter period, FS punishment was administered immediately after completion of each FR3 requirement and was thus concomitant with onset of i.v. cocaine delivery. The duration of punishment was always 0.5 s but its intensity as well as the schedule of its administration varied as a function of each specific experiment (see below).

**Figure 1:**
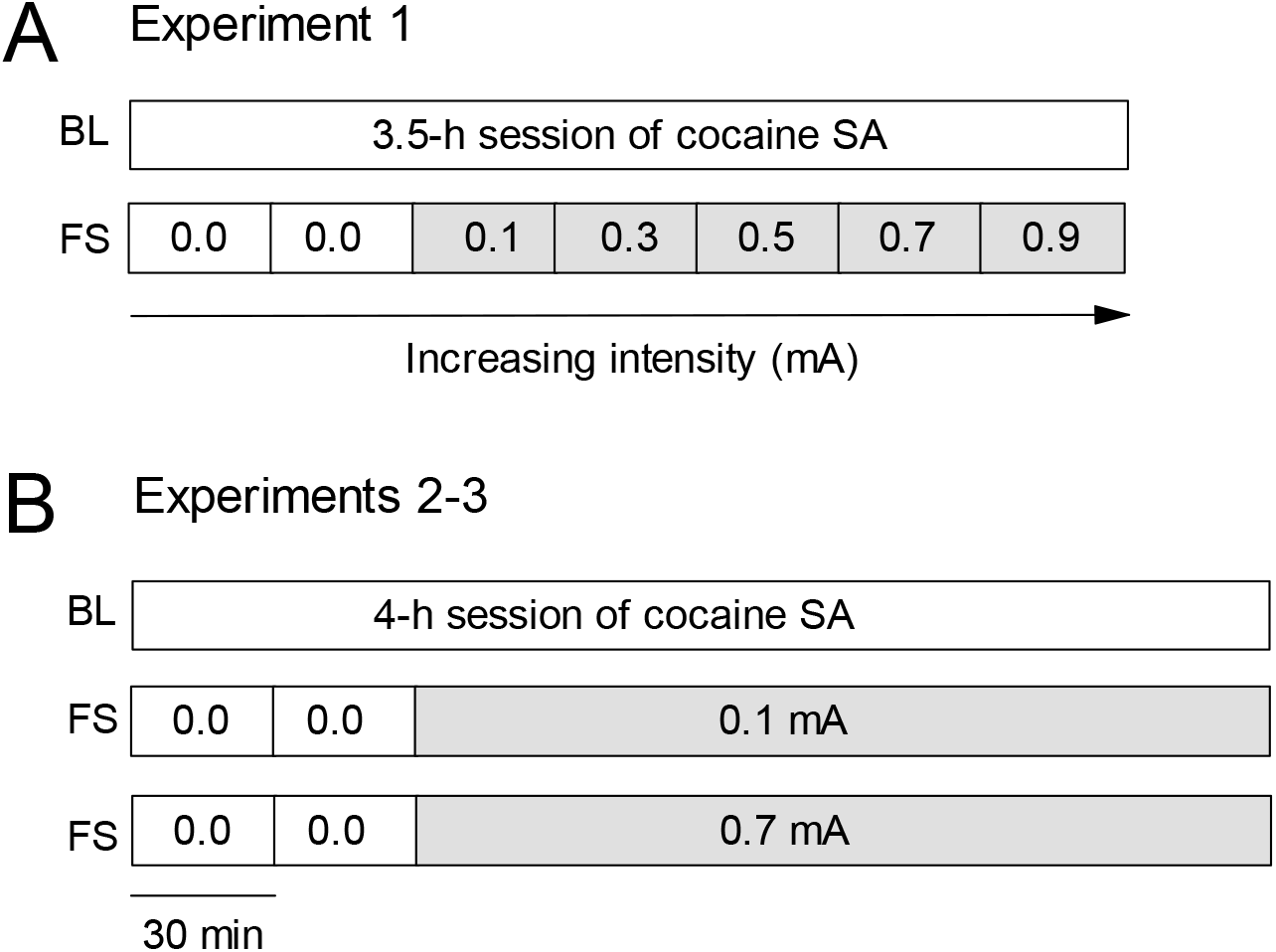
Experimental designs and procedures. (a) Measurement of resistance to footshock punishment (experiment 1). The first box represents baseline sessions (BL) with no footshock punishment of cocaine self-administration. The second box corresponds to footshock punishment sessions (FS). These sessions were subdivided into 7 30-min periods, corresponding each to a different current intensity: 0 to 0.9 mA. (b) Procedure for experiments 2 and 3. The first box represents BL sessions with no punishment. The second and third boxes represent punishment sessions with 0.1 and 0.7 mA, respectively. Note that no punishment was delivered during the first 2 30-min intervals of FS sessions.

### Specific behavioral experiments

#### Experiment 1: Measurement of resistance to footshock punishment

The goal of this initial experiment was to measure cocaine self-administration’s resistance to increasing intensity of FS punishment and its stability with repeated measurement. This experiment involved a total of 6 rats that were previously trained to self-administer cocaine during 35 3-h daily sessions. Resistance to punishment was defined operationally as the current intensity that suppressed cocaine self-administration by 50% (i.e., IS50%). To this end, cocaine self-administration was punished with ascending FS intensities from 0.1 to 0.9 mA, with an incremental step of 0.2 mA, until complete suppression of behavior. Each measurement session lasted 3.5 h and was decomposed into 7 successive 30-min intervals. As explained before, no punishment was administered during the first two 30-min intervals of cocaine self-administration (i.e., 1st hour). The following 5 30-min intervals of cocaine self-administration were each associated with a FS punishment of increasing intensity (cf. Figure 1A). To assess stability, resistance to punishment was measured two times. Importantly, between measurements, rats were retested for cocaine self-administration without punishment during several days until full recovery to baseline levels of cocaine self-administration (re-baselining). This was done to extinguish any conditioned fear to the context.

The main result of experiment 1 showed that individual resistance to punishment decreased considerably between measurements. Strikingly, as a result, rats become responsive to the lowest current intensity available that was initially behaviorally ineffective (i.e., 0.1 mA) (see Results). The following additional experiments were conducted to further characterize this acquired low resistance to FS punishment.

#### Experiment 2: Effects of prior exposure to a high-intensity FS punishment on subsequent resistance to punishment

This experiment sought to test the effects of exposure to a relatively high-intensity FS (i.e., 0.7 mA) on subsequent resistance to an initially ineffective FS intensity (i.e., 0.1 mA). This was done in a separate group of rats (n = 6) that were previously trained to self-administer cocaine during 27 2.7-h daily sessions. All daily punishment sessions lasted 4 h, with no FS during the first hour (Figure 1B). Rats were first exposed for one session to 0.1 mA. In experiment 1, this low intensity did not initially suppress cocaine self-administration in FS-naïve rats. The day after, they were exposed for one session to 0.7 mA which completely suppressed cocaine self-administration in experiment 1. After exposure to 0.7 mA, rats were retested for cocaine self-administration without punishment (i.e., re-baselining) until full recovery to pre-shock levels of drug intake. Then, they were re-exposed to the subthreshold FS intensity once a week during 5 consecutive weeks, each re-exposure occurring after every 4 sessions of re-baselining without punishment.

#### Experiment 3: Extinction of acquired low resistance to punishment

This experiment involved a separate group of rats (n = 7) that were previously trained to self-administer cocaine during 22 4-h sessions. Its design was identical to that of experiment 2, except that rats were re-exposed to 0.1-mA FS punishment during 5 consecutive daily sessions, with no intermediate re-baselining sessions. This procedure was designed to attempt to extinguish acquired low resistance to FS punishment by minimizing any spontaneous recovery process that could have taken place during the time interval between testing sessions.

#### Experiment 4: Effects of non-contingent exposure to a high-intensity FS on subsequent resistance to punishment

This experiment involved a separate group of rats (*n* = 12) that were previously trained to self-administer cocaine during 33 2.5-h sessions. Its design was identical to that of experiment 2, except that the high-intensity FS (i.e., 0.7 mA) was administered non-contingently without access to cocaine self-administration. Specifically, rats received 7 non-contingent 0.7-mA FS with variable inter-shock intervals (15-30 min) during a 4-h session. This was done in an attempt to mimic the pattern and number of response-contingent high-intensity FS received by the most exposed rats from experiments 2 and 3. After several re-baselining sessions, rats were re-exposed to 0.1-mA FS punishment during one session.

#### Experiment 5: Generalization to a different, non-painful punishment

This experiment involved a separate group of rats (*n* = 7) that were previously trained to self-administer cocaine during 19 3-h sessions. Its goal was to test whether the decreased resistance to punishment seen after FS could generalize to a different type of punishment, histamine. Histamine was shown recently to serve as an effective punisher of operant behavior reinforced by nondrug and drug rewards [41–48]. All daily histamine punishment sessions lasted 4 h, with no punishment during the first hour. During the last 3 h, cocaine self-administration was punished by co-administering histamine with cocaine upon completion of each FR3 requirement. This procedure required two syringes: one syringe containing cocaine alone and one syringe containing histamine and cocaine, the former being quickly replaced by the latter after the first hour. This was done manually in a manner to avoid introducing air in the i.v. infusion system. On the first session of punishment, rats received a low dose of histamine (i.e., 0.5 mg) with each dose of cocaine. On the following session, they received a much higher dose of histamine (i.e., 6 mg) with each dose of cocaine. These doses of histamine were selected from a pilot study. After exposure to the high dose of histamine, rats were re-tested for cocaine self-administration without histamine punishment (i.e., re-baselining) until full recovery of drug intake. Then, they were re-exposed to the low dose of histamine during one session.

## Results

### Experiment 1: Measurement of resistance to footshock punishment

Rats (n = 6) were first trained to self-administer cocaine during 35 3-h daily sessions until stabilization of drug intake. In total, they obtained 1127.5 ± 168.0 unit doses, amounting to an intake of 281.9 ± 42.0 mg. During the last 3 baseline (BL) sessions preceding punishment testing, their within-session rate of self-administration was stable during the last 6 30-min intervals at around 5 cocaine injections every 30 min (Figure 2A). The first 30-min interval of each session was systematically ignored because it includes initial drug loading (see Data Analysis). In contrast, during the first session of FS punishment (FS1), there was a current intensity-dependent suppression of cocaine self-administration compared to baseline (F5,25=27.65, p<0.01) (Figure 2A). There was initially no suppression of cocaine self-administration at 0.1 mA (the lowest current intensity tested), an intermediate level of suppression at 0.3 mA (Tuckey HSD, p<0.01) and, finally, a near complete suppression above 0.5 mA (Tuckey HSD, p<0.01). The mean current intensity that inhibits or suppresses the rate of cocaine self-administration by 50% (or IS50%; see Data Analysis) was estimated to be 0.24 ± 0.02 mA (Figure 2B). Importantly, when rats were re-tested (FS2) for FS punishment after several sessions of re-baselining, their resistance to FS punishment decreased, as indicated by a large leftward shift of the intensity-suppression curve compared to FS1 (F5,25=14.99, p<0.01) (Figure 2A) and a lowering of the IS50% (F1,5=21.43, p<0.01) (Figure 2B). As a result, rats suppressed their cocaine self-administration at 0.1 mA, the lowest FS intensity tested and to which they were initially resistant.

**Figure 2:**
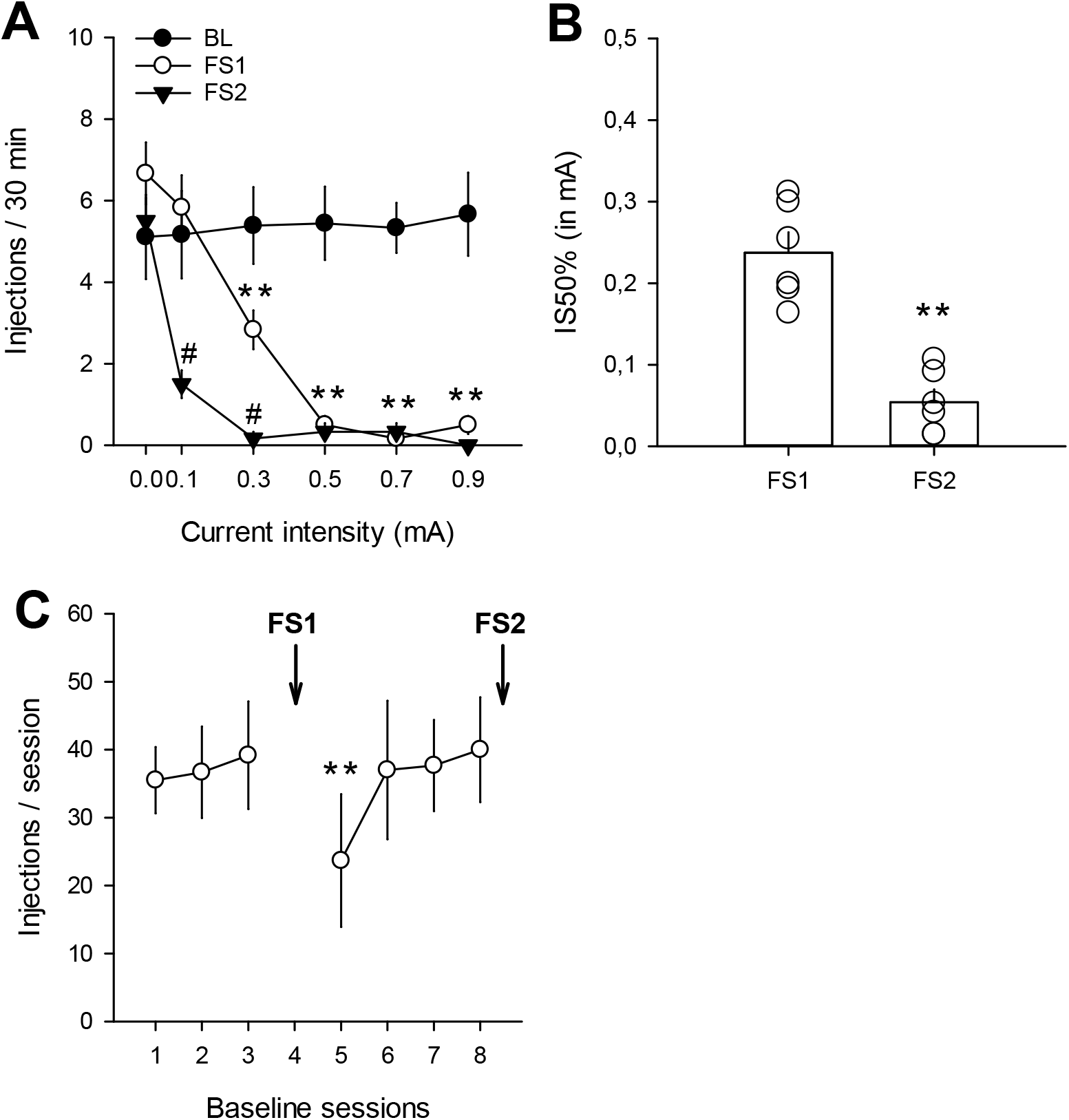
Measurement of resistance to footshock punishment. (a) Footshock punishment-induced suppression of cocaine self-administration. Number of cocaine injections (mean ± s.e.m.) during baseline (BL) or footshock punishment sessions (FS1 and FS2) as a function of increasing current intensity. ** p<0.01, different from baseline; # p<0.01, different from FS1. (b) Current intensity (mean ± s.e.m.) that suppresses by 50% (IS50%) the rate of cocaine self-administration. Lowering of IS50% during repeated exposure to FS punishment. **, different from FS1 (p < 0.01). Each point represents a different individual. (c) Progressive recovery to baseline levels of cocaine self-administration (mean ± s.e.m.) during intermediate re-baselining sessions. **, different from last baseline session before FS1 (p < 0.01).

Interestingly, during re-baselining after the first FS punishment session, there was some evidence for protracted suppression of cocaine self-administration (Figure 2C), suggesting some degree of contextual fear conditioning. However, this effect was modest (F6,30=2.59, p<0.05) and short-lived, as full recovery to initial levels of cocaine self-administration was observed as early as the second re-baselining session onward.

### Experiment 2: Effects of prior exposure to a high-intensity FS punishment on subsequent resistance to punishment

This experiment involved a separate group of 6 rats that were first trained to self-administer cocaine during 27 2.7-h daily sessions until stabilization of behavior. In total, they obtained 716.0 ± 78.0 unit doses, amounting to an intake of 179.0 ± 19.5 mg. During the last 3 baseline (BL) sessions preceding punishment testing, their within-session rate of self-administration was stable at around 5 cocaine injections every 30 min (Figure 3A, black circles), as rats from experiment 1. Then, rats were subjected on different sessions to different intensities of FS punishment (i.e., 0.1 and 0.7 mA) (see Methods). As expected, rats were initially resistant to 0.1 mA (F6,30=1.01, ns) (Figure 3A, open circles). In contrast, the day after, they almost completely stopped self-administering cocaine when the current intensity of FS punishment was increased to 0.7 mA (F6,30=7.90, p<0.01) (Figure 3A). The latter effect was relatively rapid since it began to occur during the first 30-min of punishment (i.e., 2^nd^ interval in Figure 3A). When rats were re-tested with 0.1 mA after pre-exposure to 0.7 mA and re-baselining, they stopped taking cocaine almost completely, showing a dramatic decrease in their resistance to FS punishment (F6,30=3.20, p<0.01) (Figure 3B). Though there were some fluctuations, this decreased resistance to 0.1 mA tended to persist with repeated testing between re-baselining sessions and lasted during at least one month (Punishment: F1,5=921.00, p<0.01; Punishment × Session: F4,20=0.86, ns) (Figure 3C).

**Figure 3:**
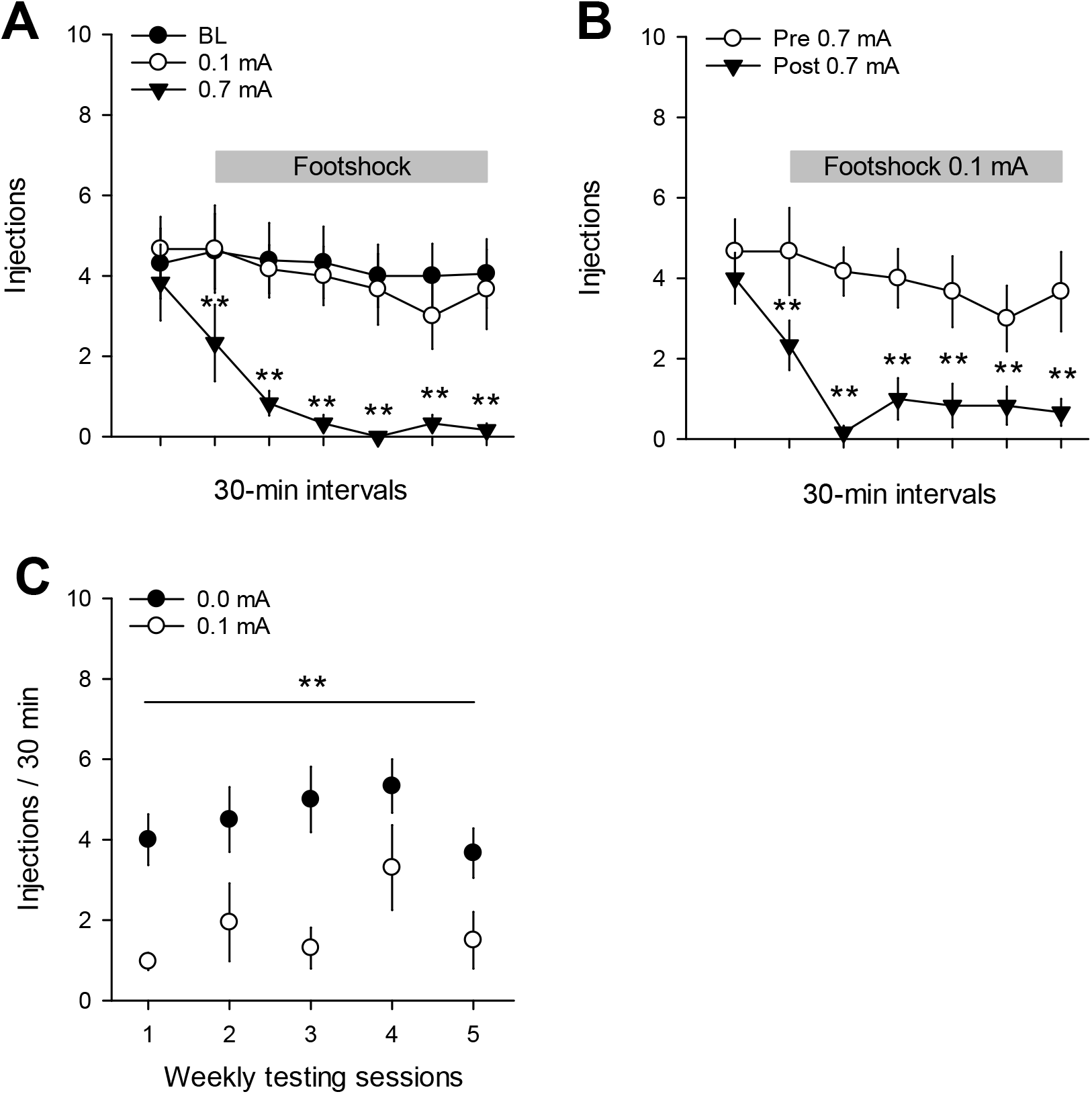
Decreased resistance to footshock punishment after exposure to high-intensity footshock punishment. (a) Number of cocaine injections (mean ± s.e.m.) during baseline sessions (BL) or footshock sessions with current intensity set to 0.1 or 0.7 mA. The horizontal grey box indicates when cocaine self-administration was punished by footshock during footshock sessions. Note that no punishment was delivered during the first 30-min interval. ** p<0.01, different from BL. (b) Effects of punishment with 0.1 mA on number of cocaine injections (mean ± s.e.m.) before and after exposure to one session with 0.7-mA footshock punishment. ** p<0.01, different from Pre 0.7 mA. (c) Persistent decrease in resistance to footshock punishment over time. Number of cocaine injections (mean ± s.e.m.) during punishment with 0.1 mA (i.e., last 6 30-min intervals) remained below control levels (i.e., 0.0 mA corresponding to the 30-min interval preceding onset of punishment) during repeated testing sessions. Testing sessions were interspersed by 4 intermediate re-baselining sessions. ** p<0.01, different from 0.0 mA.

### Experiment 3: Extinction of acquired low resistance to punishment

This experiment involved a separate group of 7 rats that were first trained to self-administer cocaine during 22 4-h daily sessions until stabilization of drug intake. In total, they obtained 928.1 ± 107.9 unit doses, amounting to an intake of 232.0 ± 27.0 mg of cocaine. During the last 3 BL sessions, their within-session rate of cocaine self-administration was stable at around 6 injections every 30 min (Figure 4A). As in experiment 2, rats were initially relatively resistant to 0.1 mA, except during the 1^st^ 30 min of exposure (F6,36=3.31, p<0.05), but lost their resistance to this intensity after having experienced one punishment session with 0.7 mA (F6,36=3.06, p<0.01) (Figure 4B) during which they nearly completely stopped to self-administer cocaine (F6,36=47.54, p<0.01) (Figure 4A). Interestingly, however, when rats were tested repeatedly with 0.1 mA during multiple consecutive sessions that were not spaced by intermediate re-baselining sessions, they rapidly recovered their initial resistance to 0.1 mA, suggesting an extinction-like effect (F4,24=2.30, p=0.08) (Figure 4C).

**Figure 4:**
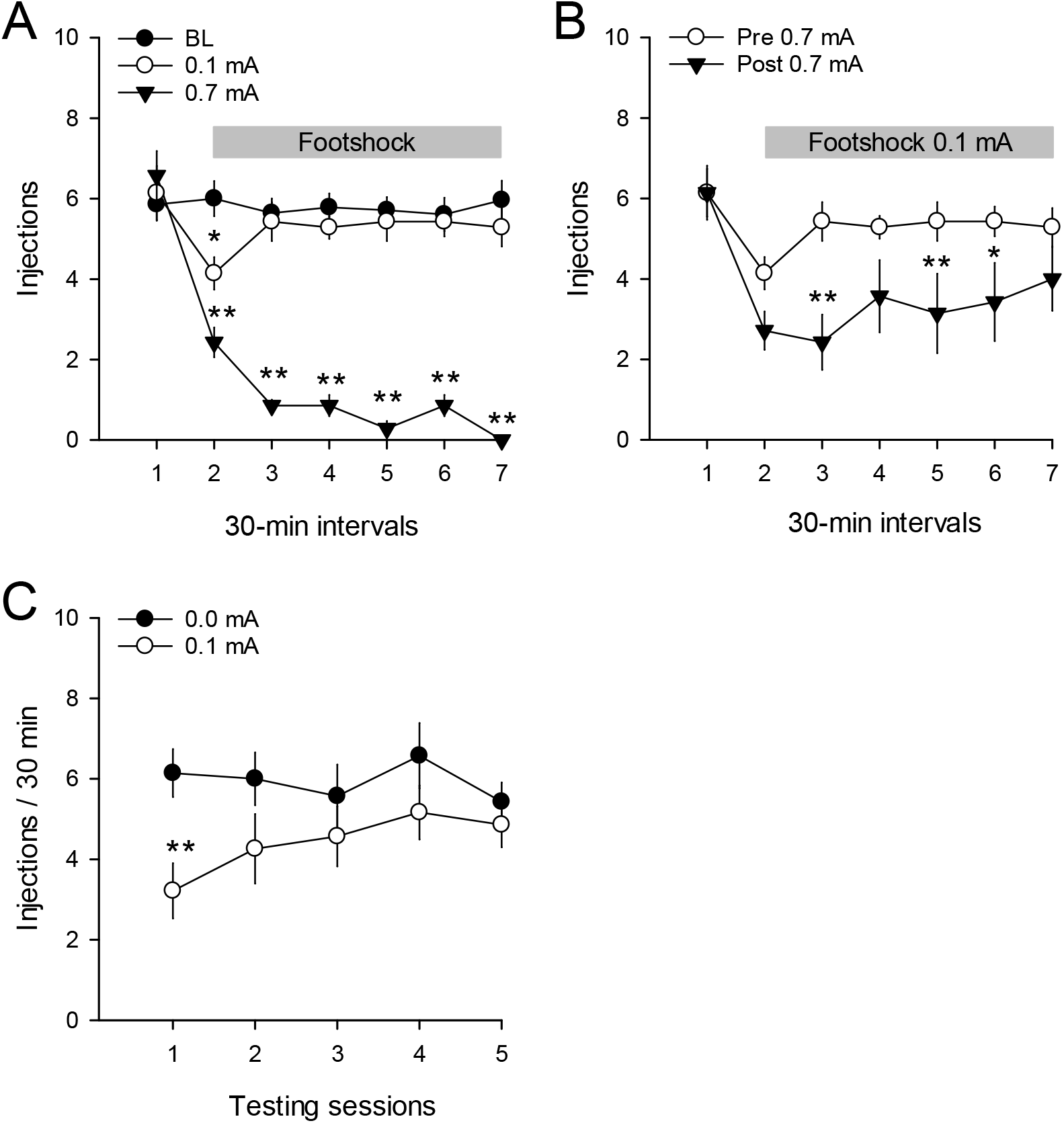
Extinction of acquired low resistance to footshock punishment. (a) Number of cocaine injections (mean ± s.e.m.) during baseline sessions (BL) or footshock sessions with current intensity set to 0.1 or 0.7 mA. * p<0.05, ** p<0.01, different from BL. (b) Effects of 0.1-mA footshock punishment on number of cocaine injections (mean ± s.e.m.) before and after exposure to 0.7-mA footshock punishment. * p<0.05, ** p<0.01, different from before exposure to 0.7 mA. (c) Extinction of acquired low resistance to footshock punishment. Number of cocaine injections (mean ± s.e.m.) during punishment with 0.1 mA (i.e., last 6 30-min intervals) compared to control levels (i.e., 0.0 mA corresponding to the 30-min interval preceding onset of punishment) during repeated testing with no intermediate re-baselining sessions. ** p<0.01, different from 0.0 mA. For additional information, see legend of Figure 3.

### Experiment 4: Effects of non-contingent exposure to a high-intensity FS on subsequent resistance to punishment

A separate group of 12 rats was used in this experiment. They were first trained to self-administer cocaine during 33 2.5-h daily sessions until stabilization of drug intake. In total, they obtained 895.7 ± 96.2 unit doses, amounting to an intake of 223.9 ± 24.0 mg of cocaine. During the last 3 BL sessions preceding punishment testing, their within-session rate of self-administration was stable at around 5 injections every 30 min (Figure 5). Rats were initially partially resistant to 0.1-mA FS punishment (Session × Interval: F6,66=2.56, p<0.05) (Figure 5A). However, in sharp contrast with previous experiments, after one session of non-contingent exposure to 0.7 mA in absence of cocaine self-administration, rats maintained their initial resistance to 0.1 mA (Session x Interval: F6,66=1.77, ns) (Figure 5B). Similar results were obtained when the analysis was confined to the subgroup of rats (n = 7) that were initially completely resistant to 0.1 mA (Figure 5C, D).

**Figure 5:**
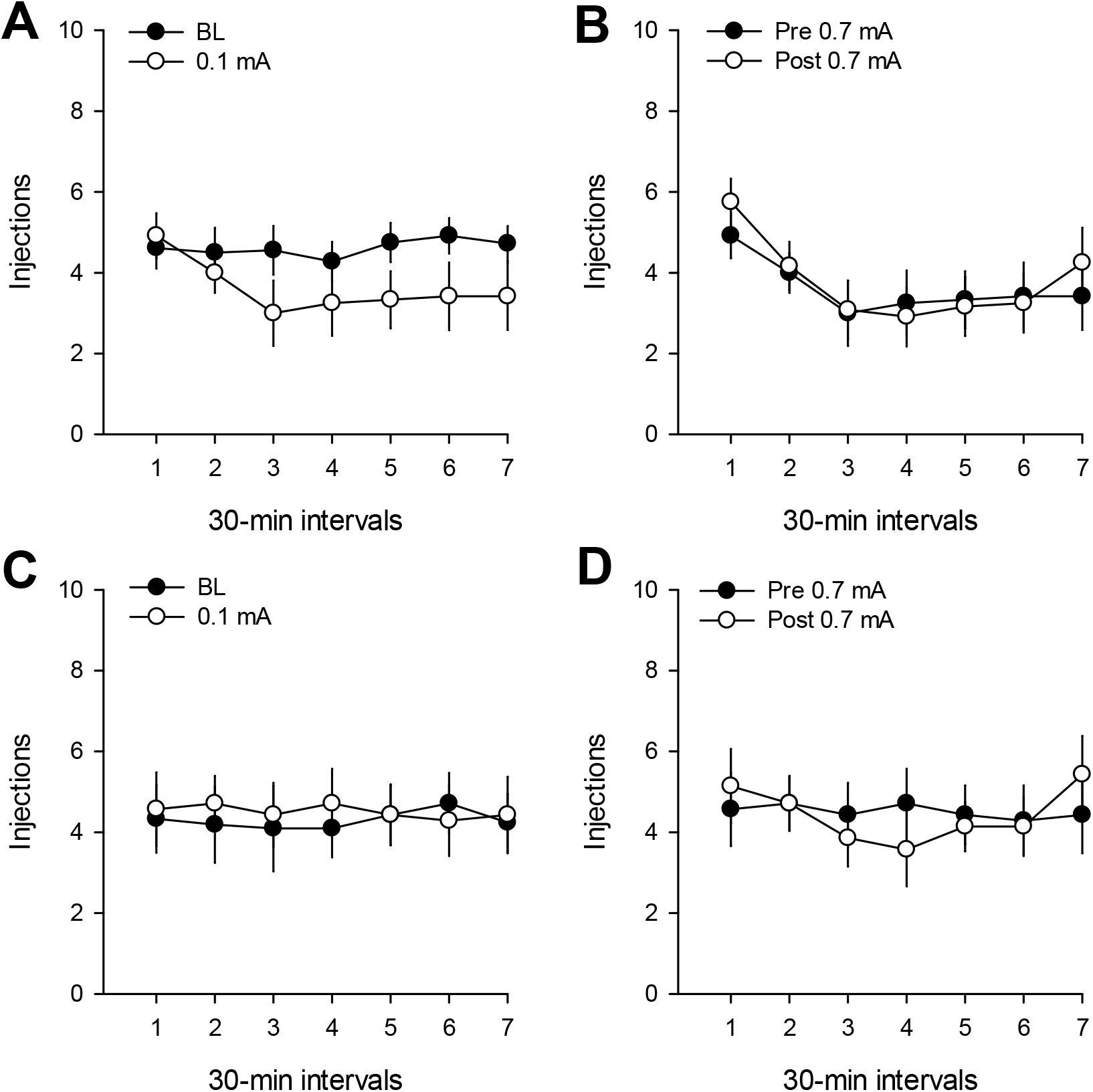
Lack of effect of non-contingent exposure to footshock. (a) Number of cocaine injections (mean ± s.e.m.) during baseline sessions (BL) or footshock sessions with current intensity set to 0.1. (b) Effects of 0.1-mA footshock punishment on number of cocaine injections (mean ± s.e.m.) before and after exposure to one session with non-contingent 0.7-mA footshock. (c), (d) idem to (a), (b), respectively, but for a subgroup of rats (n=7) that were initially insensitive to 0.1 mA.

### Experiment 5: Generalization to a different, non-painful punishment

This experiment was conducted on a separate group of 7 rats. They were first trained to self-administer cocaine during 19 3-h daily sessions until stabilization of drug intake. In total, they obtained 432.4 ± 76.2 unit doses, amounting to an intake of 108.1 ± 19.1 mg of cocaine.

During the last 3 BL sessions preceding punishment testing, their within-session rate of self-administration was stable at around 4 injections every 30 min (Figure 6A). As expected, rats did not suppress their cocaine self-administration behavior at the lowest dose of histamine (0.5 mg/injection) (F6,36=0.67, ns) but reduced considerably their intake of cocaine when their behavior was punished with the highest dose (6 mg/injection) (F6,36=2.23, p=0.06) (Figure 6A). The latter punishing effect was comparable in time-course and magnitude to that seen with 0.7 mA of footshock punishment in experiments 2 and 3. However, when rats were re-tested with the lowest dose of histamine after pre-exposure to the highest dose and re-baselining, they remained as resistant to this dose as initially (F6,36=0.67, ns) (Figure 6B).

**Figure 6:**
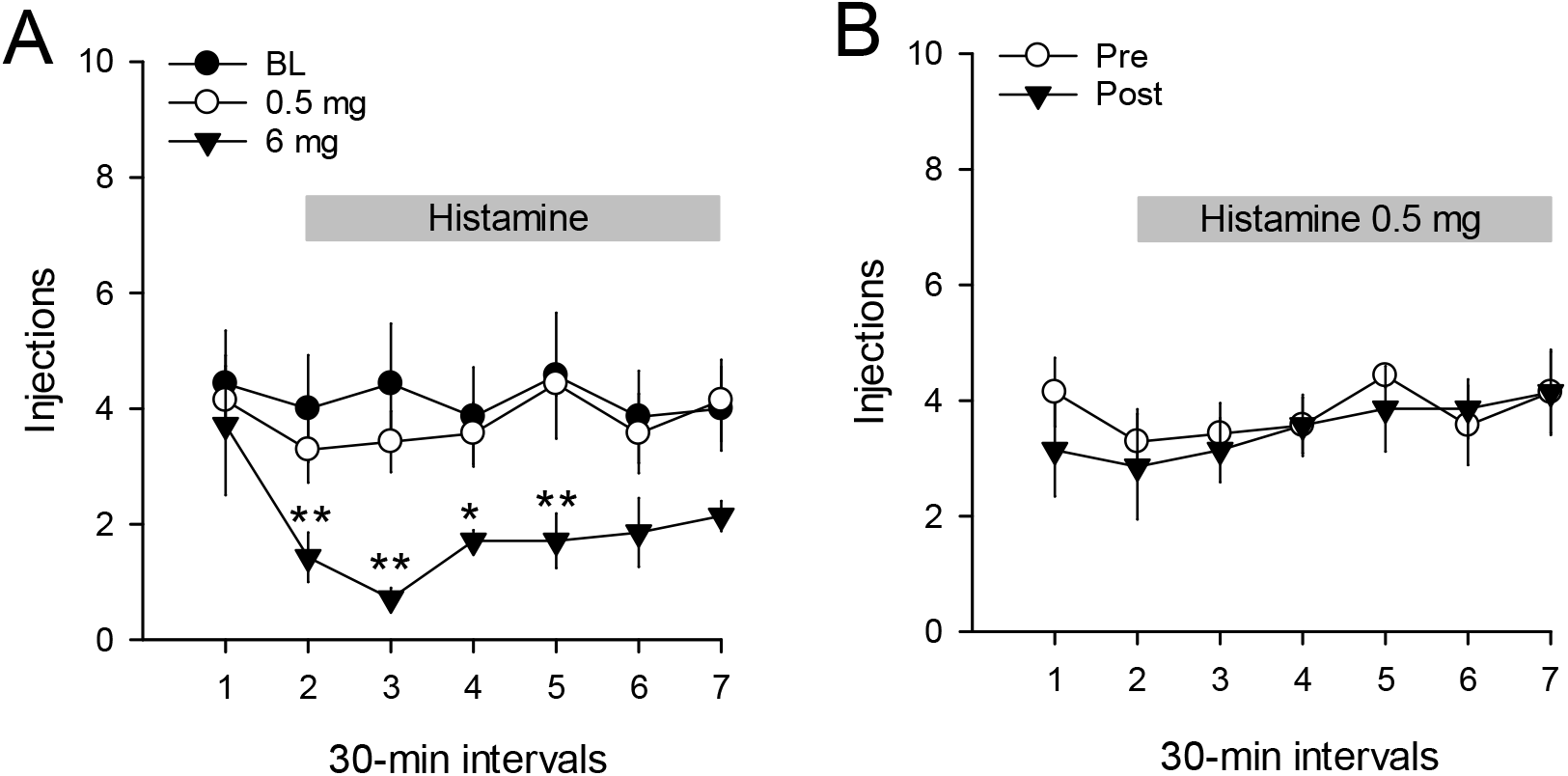
No acquired low resistance to histamine punishment. (a) Number of cocaine injections (mean ± s.e.m.) during baseline sessions (BL) or during sessions punished with 0.5 or 6 mg of i.v. histamine. The horizontal grey box indicates when cocaine self-administration was punished by histamine during corresponding punishment sessions. Note that no punishment was delivered during the first 30-min interval. * p<0.05, ** p<0.01, different from BL. (b) Effects of 0.5-mg histamine punishment on number of cocaine injections (mean ± s.e.m.) before and after exposure to one session with 6-mg histamine punishment.

## Discussion

Overall, the present study shows that cocaine self-administration’s resistance to FS punishment is not a stable individual feature, but can considerably decrease with repeated testing. Notably, we found that this acquired low resistance to punishment was mainly due to prior experience with a relatively high-intensity FS punishment that near-completely suppressed drug self-administration (i.e., 0.7 mA). After this experience, rats suppressed their cocaine intake when punished with a low-intensity and initially largely ineffective FS intensity (i.e., 0.1 mA). This decreased resistance to punishment was manifest after only few response-FS punishment pairings, but not after the same number of passive exposure to FS in the self-administration context. It occurred when the low-and high-intensity FS punishments were experienced in separate sessions and after post-punishment recovery to baseline levels of cocaine self-administration. These findings suggest that low resistance to punishment is rapidly acquired via a response-FS associative process, and does not seem to depend critically on other types of association (e.g., context-FS association). The exact nature of the underlying associative process remains to be explored [15]. Finally, this acquired low resistance to FS punishment did not generalize to a non-painful form of punishment (i.e., pharmacological punishment with histamine) that is also sometimes used to punish drug self-administration in animal studies [42,43,45,48]. The reason for this lack of generalizability is not clear at present and may suggest that acquired low resistance to punishment may be specific to painful punishment.

The phenomenon of acquired low resistance to FS punishment is not entirely new. It has been previously reported, though not discussed explicitly, in studies in which rats were re-tested with FS punishment after prior experience with a high-intensity FS punishment that near-completely suppressed drug self-administration [34,35]. Specifically, in those studies, rats were tested twice with increasing intensity of FS punishment until they showed a complete suppression of cocaine self-administration. As demonstrated here, during the second test, rats suppressed their cocaine intake when punished with FS intensities lower than during the first test. Importantly, in these studies, the intensity of FS punishment was increased between-session, not within-session as in the present study, reinforcing the generality of the observed phenomenon. Finally, it is interesting to note that studies that failed to observe an acquired low resistance to FS punishment tested rats with a relatively narrow range of intensity that only partially suppressed drug self-administration [27,36,37]. Thus, it seems that what matters most is prior experience with an intensity of FS punishment that is sufficient to near-completely suppress drug self-administration, as shown here.

However, this factor, though necessary, does not seem to be sufficient. In some studies, a range of FS intensity that completely suppressed responding for a nondrug reward did not lead to a decreased resistance to FS punishment during retesting [36,37]. Thus, it seems that acquisition of a low resistance to FS punishment is specific to cocaine self-administration, suggesting the involvement of an as-yet unidentified interaction between FS punishment and some of the pharmacological effects of cocaine. Future studies are needed to explore this possible interaction and also whether it is specific to cocaine or generalizable to other addictive drugs.

Once acquired, the persistence of this low resistance to punishment mainly depended on how often cocaine self-administering rats subsequently re-experienced low-intensity FS punishment. Acquired low resistance to punishment was extinguished within 2-3 days when low-intensity punishment was re-experienced every day but could persist during at least one month when it was re-experienced every other week. The latter observation suggests a time-dependent spontaneous recovery which is consistent with an associative account [49,50].

Time-dependent spontaneous recovery also predicts that if rats were retested one week or more after daily extinction of low resistance to FS punishment, they should also recover, at least partly, their low resistance to FS punishment. An associative account also makes several other predictions that could be tested in future research. Notably, it predicts that acquired low resistance to punishment should be context-specific and response-specific [15,50].

The present study has also implications for the interpretation of resistance of drug self-administration to FS punishment as a model of compulsion-like behavior. As explained in the Introduction, previous parametric studies, including the present study, have revealed that resistance of drug self-administration to FS punishment is intensity-dependent. When a sufficiently broad range of FS intensities is used, each individual animal eventually experiences a current intensity that leads it to stop self-administering the drug [25,28,32,34,35]. What differs between individuals is not their ability to stop drug use *per se*, but their relative resistance to FS punishment. Some individuals resist more and stop drug use at higher intensities of FS punishment than other individuals. Here we show that this relative resistance to FS punishment is not a stable individual feature as it can be lost with experience, thereby further revealing rats’ intact ability to abstain from drug use. This raises a potential dilemma for future research. If we want individual rats to keep their initial resistance to FS punishment, one must be careful not to expose them to a FS intensity that near-completely suppresses their drug self-administration behavior. However, doing that occludes the fact that rats have the ability to stop drug use and this may bias research on the mechanisms of compulsion-like behavior [13,14]. It will be interesting to know how the neurobiological mechanisms that underlie resistance to punishment at a given FS intensity change after acquisition of low resistance to punishment.

Finally, the present study has also some methodological implications. It suggests that testing drug self-administering rats with descending order of FS intensity should promote low resistance to FS punishment in comparison to ascending order of FS intensity. Indeed, a descending order of FS intensity implies exposing rats first to the highest intensity which should then reduce their resistance to subsequent lower intensities. Interestingly, previous parametric studies have typically tested drug self-administering rats with an ascending order of FS intensities [26–32,34–37,51,52]. It will be interesting to compare the effects of different order of FS intensity to determine whether and to what the acquired loss of resistance to punishment reported here can somehow be mitigated.

## Acknowledgements

We thank Christophe Bernard, Mathieu Louvet and Eric Wattelet for administrative assistance. We also thank Drs. Karine Guillem and Magalie Lenoir for their helpful comments on a previous version of the manuscript.

## Author contribution

SHA conceived the project. AD, SHA designed the experiments. AD carried out the experiments with the participation of PG and LF. AD collected the experimental data. AD, SHA analyzed the data. AD, SHA wrote the paper. All authors reviewed content and approved the final version of the manuscript.

## References

1 Association AP. Diagnostic and statistical manual of mental disorders (DSM-5). American Psychiatric Association: Washington, DC; 2013.

2 Martin CS, Langenbucher JW, Chung T, Sher KJ. Truth or consequences in the diagnosis of substance use disorders. Addiction. 2014;109(11):1773–8.

3 Hasin DS, O'Brien CP, Auriacombe M, Borges G, Bucholz K, Budney A, et al. DSM-5 criteria for substance use disorders: recommendations and rationale. Am J Psychiatry. 2013;170(8):834–51.

4 Pickard H, Ahmed SH. How do you know you have a drug problem? The role of knowledge of negative consequences in explaining drug choice in humans and rats. In: Heather N, Segal G, editors. Addiction and choice. Oxford: Oxford University Press; 2017. p. 29–48.

5 Klingemann H, Sobell MB, Sobell LC. Continuities and changes in self-change research. Addiction. 2010;105(9):1510–8.

6 Heather N. Is the concept of compulsion useful in the explanation or description of addictive behaviour and experience? Addictive Behaviors Reports. 2017;6:15–38.

7 Hogarth L. Addiction is driven by excessive goal-directed drug choice under negative affect: translational critique of habit and compulsion theory. Neuropsychopharmacology. 2020;45(5):720–35.

8 Heyman GM. Quitting drugs: quantitative and qualitative features. Annu Rev Clin Psychol. 2013;9:29–59.

9 Smith GS, Davis M. Punishment of amphetamine and morphine self-administration behavior. The psychological record. 1974;24:477–80.

10 Grove RN, Schuster CR. Suppression of cocaine self-administration by extinction and punishment. Pharmacol Biochem Behav. 1974;2(2):199–208.

11 Bergman J, Johanson CE. The effects of electric shock on responding maintained by cocaine in rhesus monkeys. Pharmacol Biochem Behav. 1981;14(3):423–6.

12 Wolffgramm J, Heyne A. From controlled drug intake to loss of control: the irreversible development of drug addiction in the rat. Behav Brain Res. 1995;70(1):77–94.

13 Vanderschuren LJMJ, Minnaard AM, Smeets JAS, Lesscher HM. Punishment models of addictive behavior. Current Opinion in Behavioral Sciences. 2017;13:77–84.

14 Vanderschuren L, Ahmed SH. Animal Models of the Behavioral Symptoms of Substance Use Disorders. Cold Spring Harbor perspectives in medicine. 2020.

15 Jean-Richard-Dit-Bressel P, Killcross S, McNally GP. Behavioral and neurobiological mechanisms of punishment: implications for psychiatric disorders. Neuropsychopharmacology. 2018;43(8):1639–50.

16 Ahmed SH. “A walk on the wild side” of addiction: the history and significance of animal models. In: Pickard H, Ahmed SH, editors. The Routledge Handbook of Philosophy and Science of Addiction. New York: Routledge; 2019. p. 192–204.

17 Deroche-Gamonet V, Belin D, Piazza PV. Evidence for addiction-like behavior in the rat. Science. 2004;305(5686):1014–7.

18 Panlilio LV, Thorndike EB, Schindler CW. Reinstatement of punishment-suppressed opioid self-administration in rats: an alternative model of relapse to drug abuse. Psychopharmacology (Berl). 2003;168(1–2):229–35.

19 Chen BT, Yau HJ, Hatch C, Kusumoto-Yoshida I, Cho SL, Hopf FW, et al. Rescuing cocaine-induced prefrontal cortex hypoactivity prevents compulsive cocaine seeking. Nature. 2013;496(7445):359–62.

20 Pascoli V, Terrier J, Hiver A, Luscher C. Sufficiency of Mesolimbic Dopamine Neuron Stimulation for the Progression to Addiction. Neuron. 2015;88(5):1054–66.

21 Ahmed SH. Validation crisis in animal models of drug addiction: beyond non-disordered drug use toward drug addiction. Neurosci Biobehav Rev. 2010;35(2):172–84.

22 Ahmed SH. The science of making drug-addicted animals. Neuroscience. 2012;211:107–25.

23 Kasanetz F, Deroche-Gamonet V, Berson N, Balado E, Lafourcade M, Manzoni O, et al. Transition to addiction is associated with a persistent impairment in synaptic plasticity. Science. 2010;328(5986):1709–12.

24 Kasanetz F, Lafourcade M, Deroche-Gamonet V, Revest JM, Berson N, Balado E, et al. Prefrontal synaptic markers of cocaine addiction-like behavior in rats. Mol Psychiatry. 2013;18(6):729–37.

25 Farrell MR, Ruiz CM, Castillo E, Faget L, Khanbijian C, Liu S, et al. Ventral pallidum is essential for cocaine relapse after voluntary abstinence in rats. Neuropsychopharmacology. 2019;44(13):2174–85.

26 Giuliano C, Belin D, Everitt BJ. Compulsive Alcohol Seeking Results from a Failure to Disengage Dorsolateral Striatal Control over Behavior. J Neurosci. 2019;39(9):1744–54.

27 Giuliano C, Pena-Oliver Y, Goodlett CR, Cardinal RN, Robbins TW, Bullmore ET, et al. Evidence for a Long-Lasting Compulsive Alcohol Seeking Phenotype in Rats. Neuropsychopharmacology. 2018;43(4):728–38.

28 Krasnova IN, Marchant NJ, Ladenheim B, McCoy MT, Panlilio LV, Bossert JM, et al. Incubation of methamphetamine and palatable food craving after punishment-induced abstinence. Neuropsychopharmacology. 2014;39(8):2008–16.

29 Torres OV, Jayanthi S, Ladenheim B, McCoy MT, Krasnova IN, Cadet JL. Compulsive methamphetamine taking under punishment is associated with greater cue-induced drug seeking in rats. Behav Brain Res. 2017;326:265–71.

30 Marchant NJ, Campbell EJ, Whitaker LR, Harvey BK, Kaganovsky K, Adhikary S, et al. Role of Ventral Subiculum in Context-Induced Relapse to Alcohol Seeking after Punishment-Imposed Abstinence. J Neurosci. 2016;36(11):3281–94.

31 Marchant NJ, Kaganovsky K, Shaham Y, Bossert JM. Role of corticostriatal circuits in context-induced reinstatement of drug seeking. Brain Res. 2015;1628(Pt A):219–32.

32 Marchant NJ, Khuc TN, Pickens CL, Bonci A, Shaham Y. Context-induced relapse to alcohol seeking after punishment in a rat model. Biol Psychiatry. 2013;73(3):256–62.

33 Marchant NJ, Rabei R, Kaganovsky K, Caprioli D, Bossert JM, Bonci A, et al. A critical role of lateral hypothalamus in context-induced relapse to alcohol seeking after punishment-imposed abstinence. J Neurosci. 2014;34(22):7447–57.

34 Pelloux Y, Hoots JK, Cifani C, Adhikary S, Martin J, Minier-Toribio A, et al. Context-induced relapse to cocaine seeking after punishment-imposed abstinence is associated with activation of cortical and subcortical brain regions. Addict Biol. 2018;23(2):699–712.

35 Pelloux Y, Minier-Toribio A, Hoots JK, Bossert JM, Shaham Y. Opposite Effects of Basolateral Amygdala Inactivation on Context-Induced Relapse to Cocaine Seeking after Extinction versus Punishment. J Neurosci. 2018;38(1):51–59.

36 Datta U, Martini M, Fan M, Sun W. Compulsive sucrose- and cocaine-seeking behaviors in male and female Wistar rats. Psychopharmacology (Berl). 2018;235(8):2395–405.

37 Datta U, Martini M, Sun W. Different functional domains measured by cocaine self-administration under the progressive-ratio and punishment schedules in male Wistar rats. Psychopharmacology (Berl). 2018;235(3):897–907.

38 Ahmed SH, Koob GF. Transition to drug addiction: a negative reinforcement model based on an allostatic decrease in reward function. Psychopharmacology (Berl). 2005;180(3):473–90.

39 Pickens R, Meisch RA, Thompson T. Drug self-administration: an analysis of the reinforcing effects of drugs. In: Iversen LL, Iversen S.D., Solomon S.H., editor Handbook of Psychopharmacology: Drugs of Abuse. New York: Plenum Press; 1978. p. 1–37.

40 Yokel RA. Intravenous self-administration: response rates, the effects of pharmacological challenges, and drug preference. In: Bozarth MA, editor Methods of Assessing the Reinforcing Properties of Abused Drugs. New York: Springer-Verlag; 1987. p. 1–33.

41 Holtz NA, Carroll ME. Cocaine self-administration punished by intravenous histamine in adolescent and adult rats. Behav Pharmacol. 2015;26(4):393–7.

42 Negus SS. Effects of punishment on choice between cocaine and food in rhesus monkeys. Psychopharmacology (Berl). 2005;181(2):244–52.

43 Gancarz-Kausch AM, Adank DN, Dietz DM. Prolonged withdrawal following cocaine self-administration increases resistance to punishment in a cocaine binge. Scientific reports. 2014;4:6876.

44 Goldberg SR. Histamine as a punisher in squirrel monkeys: effects of pentobarbital, chlordiazepoxide and H1- and H2-receptor antagonists on behavior and cardiovascular responses. J Pharmacol Exp Ther. 1980;214(3):726–36.

45 Holtz NA, Anker JJ, Regier PS, Claxton A, Carroll ME. Cocaine self-administration punished by i.v. histamine in rat models of high and low drug abuse vulnerability: effects of saccharin preference, impulsivity, and sex. Physiol Behav. 2013;122:32–8.

46 Podlesnik CA, Jimenez-Gomez C, Woods JH. A choice procedure to assess the aversive effects of drugs in rodents. J Exp Anal Behav. 2010;93(2):203–23.

47 Woolverton WL. A novel choice method for studying drugs as punishers. Pharmacol Biochem Behav. 2003;76(1):125–31.

48 Woolverton WL, Freeman KB, Myerson J, Green L. Suppression of cocaine self-administration in monkeys: effects of delayed punishment. Psychopharmacology (Berl). 2012;220(3):509–17.

49 Delamater AR, Westbrook RF. Psychological and neural mechanisms of experimental extinction: a selective review. Neurobiology of learning and memory. 2014;108:38–51.

50 Bouton ME. Extinction of instrumental (operant) learning: interference, varieties of context, and mechanisms of contextual control. Psychopharmacology (Berl). 2019;236(1):7–19.

51 Krasnova IN, Gerra MC, Walther D, Jayanthi S, Ladenheim B, McCoy MT, et al. Compulsive methamphetamine taking in the presence of punishment is associated with increased oxytocin expression in the nucleus accumbens of rats. Scientific reports. 2017;7(1):8331.

52 Torres OV, Jayanthi S, McCoy MT, Cadet JL. Selective Activation of Striatal NGF-TrkA/p75NTR/MAPK Intracellular Signaling in Rats That Show Suppression of Methamphetamine Intake 30 Days following Drug Abstinence. The international journal of neuropsychopharmacology. 2018;21(3):281–90.

